# Evaluating the evidence for biotypes of depression: attempted replication of Drysdale et.al. 2017

**DOI:** 10.1101/416321

**Authors:** Richard Dinga, Lianne Schmaal, Brenda Penninx, Marie Jose van Tol, Dick J. Veltman, Laura van Velzen, Nic van der Wee, Andre Marquand

## Abstract

**Background:** Psychiatric disorders are highly heterogeneous, defined based on symptoms with little connection to potential underlying biological mechanisms. A possible approach to dissect biological heterogeneity is to look for biologically meaningful subtypes. A recent study Drysdale et al. (2017) showed promising results along this line by simultaneously using resting state fMRI and clinical data and identified four distinct subtypes of depression with different clinical profiles and abnormal resting state fMRI connectivity. These subtypes were predictive of treatment response to transcranial magnetic stimulation therapy.

**Objective:** Here, we attempted to replicate the procedure followed in the Drysdale et al. study and their findings in an independent dataset of a clinically more heterogeneous sample of 187 participants with depression and anxiety. We aimed to answer the following questions: 1) Using the same procedure, can we find a statistically significant and reliable relationship between brain connectivity and clinical symptoms? 2) Is the observed relationship similar to the one found in the original study? 3) Can we identify distinct and reliable subtypes? 4) Do they have similar clinical profiles as the subtypes identified in the original study?

**Methods:** We followed the original procedure as closely as possible, including a canonical correlation analysis to find a low dimensional representation of clinically relevant resting state fMRI features, followed by hierarchical clustering to identify subtypes. We extended the original <u>procedure</u> using additional statistical tests, to test the statistical significance of the relationship between resting state fMRI and clinical data, and the existence of distinct subtypes. Furthermore, we examined the stability of the whole procedure using resampling.

**Results and Conclusion:** We were not able to replicate the findings of the original study. Relationships between brain connectivity and clinical symptoms were not statistically significant and we also did not find clearly distinct subtypes of depression. We argue, that based on our rigorous approach and in-depth review of the original results, that the evidence for the existence of the distinct resting state connectivity based subtypes of depression is weak and should be interpreted with caution.

## 1. INTRODUCTION

Psychiatric disorders are highly heterogeneous in terms of symptom presentation and underlying biological mechanisms and are diagnosed exclusively in terms of symptoms, which may not correspond to biological causes [2]. This, together with frequent comorbidities between disorders, complicates clinical diagnosis and hinders efforts to understand biological mechanisms of disorders and to develop better treatments. This problem has been known for a long time, but little progress has been made and clinical decision-making is still mostly done on the basis of symptoms. Recent initiatives such as the Research Domain Criteria (RDoC) [3] aim to address this issue of heterogeneity by going beyond current diagnostic categories and focusing analysis on different domains of functioning and pathology across multiple levels of analysis, including clinical symptoms, behavior, and biology.

Many studies have used data-driven clustering methods in order to find new subgroups of clinical populations, based on either clinical or biological data, with some degree of success [4]. However, these putative subgroups generally show moderate to poor reproducibility across studies, have not been extensively validated against clinical outcomes and as a consequence still have not translated into clinical practice. For example, the dominant approach of clustering based on clinical symptoms alone can provide new insights into psychopathology, however, it may not yield subtypes that reflect underlying biological differences. On the other hand, the variability of biological data is more often than not unrelated to any specific psychiatric disorder or symptom class. Thus clustering based on biological data alone may detect subtypes that are unrelated to psychiatric pathologies, and instead reflect dominant nuisance variance in the data such as groups of people with similar brain size or body type or common ancestry in the case of genetics. One way to overcome these limitations is to constrain the search for subtypes in biological data to lie along axes of variance that are related to psychiatric symptomatology. However few studies have used such an approach [4].

A prominent example following this approach is a recent study by Drysdale and colleagues [1] that aimed to stratify major depressive disorder (MDD) on the basis of biology and behavior and suggested the existence of four distinct ‘biotypes’. The authors used canonical correlation analysis (CCA) [5] to identify a two-dimensional mapping between functional connectivity measures derived from resting state fMRI (RS-fMRI) data and MDD symptoms. CCA is a well-established method for finding multivariate associations between different data sources and has been used extensively in clinical neuroimaging, for example for finding associations between neuroimaging data and behavior [6,7] and neuroimaging and genetics [8]. Next, Drysdale et al. applied a hierarchical clustering on two components derived from CCA and identified four different clusters of MDD patients, i.e. the aforementioned ‘biotypes’. Impressively, these biotypes were predictive of transcranial magnetic stimulation (TMS) treatment response, and they were also evaluated in an independent sample. However, the study has some methodological limitations. For example, the existence of distinct clusters was not conclusively established in that the authors did not test the possibility that subjects were sampled from a single continuous distribution without underlying clusters. Nevertheless, the results are promising, and if replicated it would be an important step towards understanding biological mechanisms of MDD.

The aim of this study is to replicate the biotypes identified by Drysdale et al. in a completely independent sample, namely data from the Netherlands Study of Depression and Anxiety (NESDA) [9] and Mood Treatment with Antidepressants or Running (MOTAR) study (Lever-van Milligen et al., in preparation). These studies together create a relatively large cohort containing a heterogeneous naturalistic sample of subjects with depression, anxiety and depression-anxiety comorbidity, thus capturing a wider range of possible clinical and biological profiles relative to the study by Drysdale and colleagues, which included mainly hospitalized treatment-resistant patients. The original study used 220 patients as a cluster discovery dataset and an additional 92 patients from the same cohort as a replication dataset. Our combined dataset of NESDA and MOTAR includes a cohort of 187 participants with comparable clinical measures to the original study. Specifically, we aimed to answer the following questions: 1) Using the same procedure as in [1], can we find a statistically significant and reliable relationship between brain connectivity and clinical symptoms? 2) Is the identified relationship similar to the one found in the original study? 3) Can we identify distinct and reliable subtypes? 4) If so, do they have similar clinical profiles as the subtypes identified in the original study? Subsequently, we will also perform a critical evaluation of methods used by Drysdale et al. and provide a recommendation for future studies.

## 2. METHODS

We conducted our analysis as close to possible and to our best understanding of the published analysis pipeline in the Drysdale et al. study. Several details related to the analysis were not specified in the original paper and were clarified via personal communication with the corresponding author. We included several additional validation steps for CCA and cluster analysis. Our aim was to replicate the analysis steps related to the creation and evaluation of subtypes, we did not try to replicate additional analyses performed in the original study such as classification of healthy and depressed subjects or the prediction of TMS treatment response. We describe our pipeline and the Drysdale et al. pipeline below and provide a side by side schematic comparison in Figure 1.

**Figure 1:**
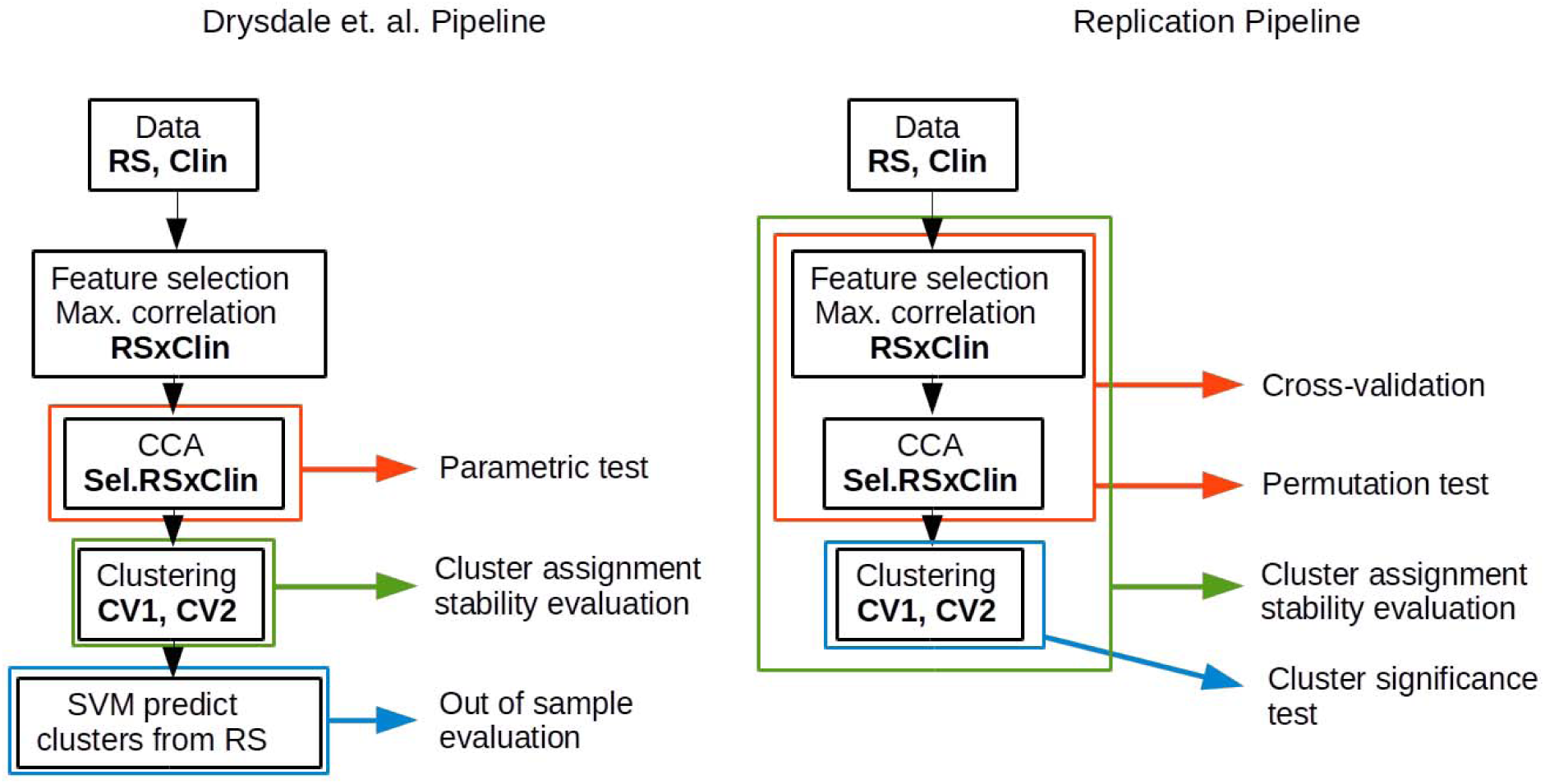
A scheme of pipeline used in the original study and our pipeline. **Data**: in the original study, 220 depressed subjects have been analyzed as a part of a “cluster discovery” set an additional 92 subjects as evaluation set. The clinical data (**Clin**) consisted of 17 HAM-D items. We have used 187 subjects with depression, anxiety or depression-anxiety comorbidity. The clinical data consisted of 17 IDS items, that best-matched HAM-D item used in the original study. After preprocessing of fMRI data (**RS**), a correlation matrix between selected regions had been created, resulting in ∼35000 features. A small subset of features (178 in the original study and 150 in our study) was selected based on their correlation with clinical symptoms (**Sel.RS**). Then, CCA was performed using these selected features and clinical symptoms. In the original study, a parametric test was used to the established statistical significance of CCA without taking a previous feature selection into an account. Hierarchical clustering was performed on first two resting state connectivity canonical variates (**CV1, CV2**). We have included additional test, to test if the data cluster more than what is expected from data sampled from a Gaussian distribution. Stability of cluster assignment was evaluated in the original study by resampling of CV1 and CV2, we have evaluated the stability under resampling but by repeating also feature selection and CCA procedures. **Out of sample evaluation**: in the original study, additional 92 subjects have been assigned to clusters according to SVM model and clinical profiles of these clusters have been compared to clinical profiles of clusters obtained in the cluster discovery set. We have evaluated the reproducibility of canonical correlations directly, using 10-fold cross-validation.

### 2.1 Sample characteristics

All our analyses were performed on 187 subjects from NESDA and MOTAR samples diagnosed according to DSM-IV criteria with MDD, anxiety (panic, social phobia or generalized anxiety disorder) or both MDD and anxiety, established using the structured Composite International Diagnostic Interview (CIDI, version 2.1) [10].

The original Drysdale et al. study included 220 subjects in a cluster discovery set and an additional 92 subjects in a validation set with an active episode of MDD and a history of treatment resistance. In our sample, 151 subjects came from the baseline assessment of NESDA [9], which is large naturalistic cohort study of depression and anxiety. Additional descriptions of the NESDA cohort can be found in [11]. An additional 36 subjects were from the baseline assessment of the MOTAR study (Lever-van Milligen et al., in preparation), which is a randomized controlled treatment study (antidepressants or running therapy).

All participants gave written informed consent. Studies were approved by the Central Ethics Committees of the participating medical centers: Leiden University Medical Center (LUMC), Amsterdam Medical Center (AMC), and University Medical Center Groningen (UMCG) for NESDA and Ethics committee of Amsterdam Medical Center (AMC) for MOTAR.

### 2.2 Resting-state fMRI

Participants from NESDA were scanned at one of the three participating scan centers and at one scan center for the MOTAR study. All imaging data were acquired on a Philips 3.0-T Achieva MRI scanner. RS-fMRI data were acquired using T2*-weighted gradient-echo echo-planar imaging with the following scan parameters for the NESDA sample: Amsterdam and Leiden centers: 200 whole-brain volumes; repetition time (TR) = 2300 ms; echo time (TE) = 30 ms; flip angle = 80º; 35 axial slices; no slice gap; FOV = 220 × 220 mm; in plane voxel resolution = 2.3 mm × 2.3 mm; slice thickness = 3 mm; same in Groningen, except: TE = 28 ms; 39 axial slices; in plane voxel resolution = 3.45 mm × 3.45 mm. And for the MOTAR sample: 210 whole-brain volumes; repetition time (TR) = 2300 ms; echo time (TE) = 28 ms, flip angle= 76.1º, 37 axial slices, no slice gap; FOV = 240×240, in plane voxel resolution = 3.3 mm × 3.3 mm. T1-weighted image was acquired with the repetition time (TR) = 9 ms; echo time (TE) = 3.5 ms; flip angle = 8º; 170 sagittal slices; no slice gap; FOV = 256 × 256 mm; in plane voxel resolution = 1 mm × 1 mm; slice thickness = 1 mm.

Preprocessing of RS-fMRI data was performed using FSL 5.0.8 and included motion correction, grand mean scaling of the fMRI time series, spatial smoothing with 6mm Gaussian kernel, motion artefacts removal using ICA-AROMA [12], nuisance signal regression of white matter and CSF, and high pass filtering with a cut-off frequency of 100 seconds. The resulting RS-fMRI images were registered to Montreal Neurological Institute (MNI) space using registration matrices obtained from the first co-registration of functional images to T1 image using the boundary based registration tool and registering the T1 images to MNI template brain.

Next, correlation matrices were created using 264 cortical parcellations proposed by Power et al. [13] plus an additional 13 regions, including the left and right caudate, amygdala, hippocampus, nucleus accumbens and subgenual anterior cingulate cortex, as in the original Drysdale et al. study. We averaged all voxels within each region to create a single time series per region and then we created a correlation matrix by computing pairwise Pearson’s correlation coefficients between all regions. This resulted in 38,000 connectivity features (lower diagonal of 277*277 correlation matrix), which were later reduced to 37,675 by discarding regions with insufficient coverage in more than 10% of subjects. These correlations were transformed using Fisher’s z-transform and linear effects of age and scan location were regressed out.

### 2.3 Clinical characteristics

The original study used depressive symptom scores of the Hamilton rating scale for depression (HAMD) [14] in their analyses. Here we used depressive symptom scores derived from the Inventory of Depressive Symptomatology (IDS) [15]. This inventory was developed as an improvement over HAMD, aiming to improve the coverage of common MDD symptoms [16]. However, to make our study comparable to the original study, we used only a subset of 17 IDS items that best matched the items of the HAMD (Table 1).

**Table 1:**
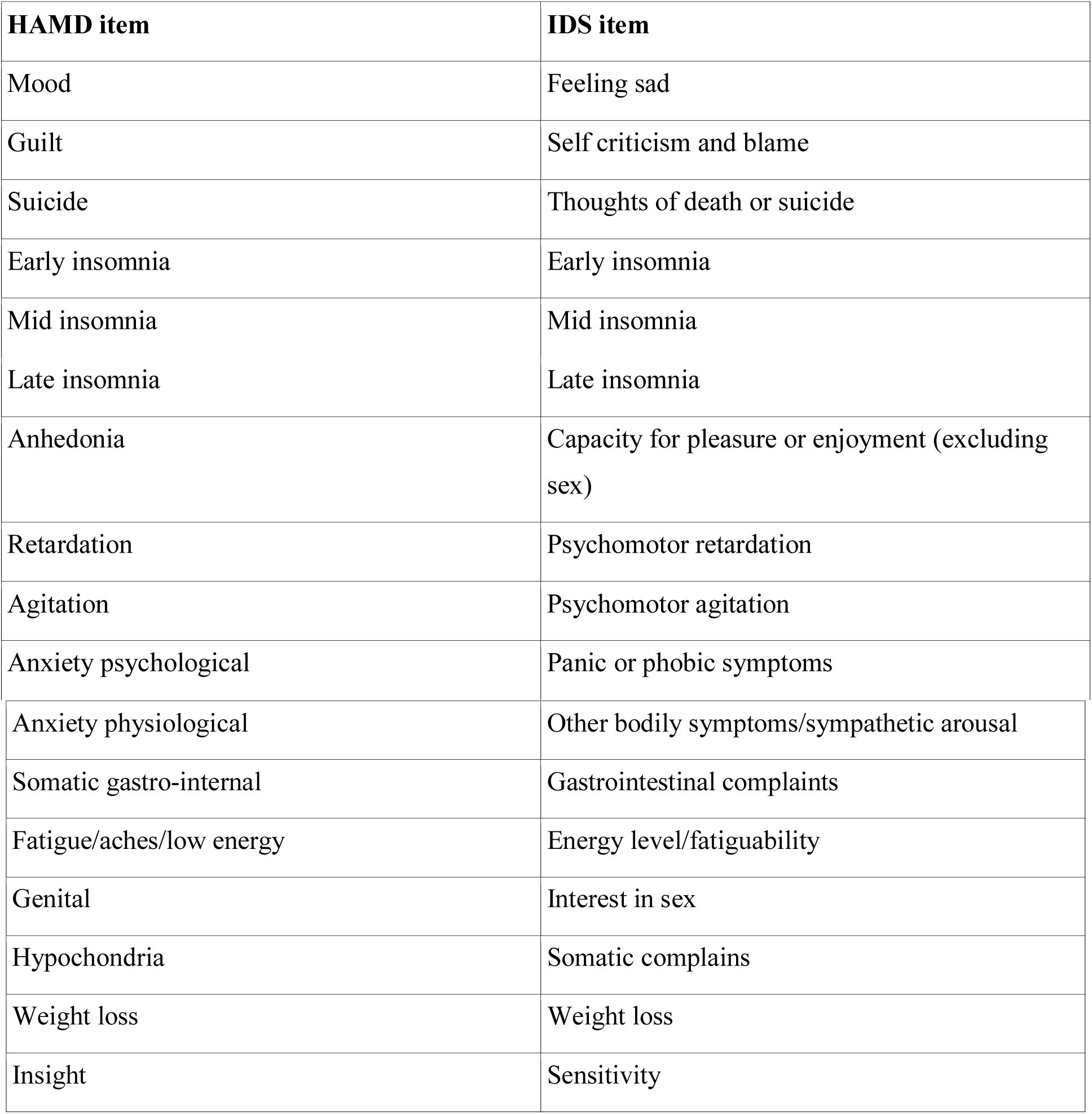
HAM-D items used in the original study and best-matched IDS items used in this study.

| HAMD item | IDS item |
| --- | --- |
| Mood | Feeling sad |
| Guilt | Self criticism and blame |
| Suicide | Thoughts of death or suicide |
| Early insomnia | Early insomnia |
| Mid insomnia | Mid insomnia |
| Late insomnia | Late insomnia |
| Anhedonia | Capacity for pleasure or enjoyment (excluding sex) |
| Retardation | Psychomotor retardation |
| Agitation | Psychomotor agitation |
| Anxiety psychological | Panic or phobic symptoms |
| Anxiety physiological | Other bodily symptoms/sympathetic arousal |
| Somatic gastro-intestinal | Gastrointestinal complaints |
| Fatigue/aches/low energy | Energy level/fatiguability |
| Genital | Interest in sex |
| Hypochondria | Somatic complains |
| Weight loss | Weight loss |
| Insight | Sensitivity |

### 2.4 Feature selection

From the 37,675 connectivity features (the lower diagonal of the functional correlation matrix), we selected the subset of features with the highest Spearman’s correlation with any of the 17 IDS symptoms. In the original study, according to communication with the corresponding author, during feature selection, the number of RS-fMRI features corresponding to approximately 80% of the total number of participants were retained, which corresponds to 176 RS-fMRI connectivity features in a sample of 220 participants in the original study. Here we selected the top 150 RS-fMRI features with the highest Spearman’s correlation with any of the 17 IDS symptoms to preserve the same feature to subjects ratio (80% of 187 subjects).

### 2.5 Canonical correlation analysis

Next, following the original study, we performed a canonical correlation analysis (CCA) on the selected RS-fMRI connectivity features and depressive symptoms. Canonical correlation analysis [5] is a multivariate statistical method that seeks an association between two sets of variables. CCA is the most general multivariate technique with multiple regression, MANOVA, and discriminant analysis all as special cases of CCA (multiple regression is a CCA with only one variable in Y, MANOVA, and discriminant analysis are CCA with binary variables in X or Y). Given the two multidimensional datasets X (e.g. clinical features) and Y (e.g. RS-fMRI connectivity features), canonical correlation analysis finds a linear combination of X that maximally correlates with a linear combination of Y. This linear combinations of X and Y are new variables, called canonical variates. Both canonical variates for X and Y are called a canonical pair and the correlation between canonical variates is called a canonical correlation. Multiple canonical pairs can be found with a constraint that each subsequent canonical pair has to be uncorrelated with all the previous ones.

CCA is also closely related to PCA with a difference that CCA performs eigen decomposition of the cross-correlation matrix instead of the correlation matrix. In PCA the first principal component explains the largest amount of variance in the data, and each subsequent principal component explains a (smaller) maximal amount of variance that is orthogonal to all the previous ones. In CCA the first canonical variate of X explains the largest amount of variance in Y and each subsequent canonical variate is explaining less of variance in Y and is orthogonal to all the previous canonical variates. In more detail, the squared canonical correlations are eigenvalues of the matrix:

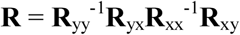

where **R**_xx_ and **R**_yy_ are correlation matrices of X and Y, respectively, and **R**_xy_ and **R**_yx_ are cross-correlation matrices of variables from X with variables from Y. Coefficients that create the canonical variates are the respective eigenvectors of R. CCA can be also thought of as a dimensionality reduction step, where the original data of X and Y are mapped into a lower dimensional space of canonical variates whose dimensions are highly correlated between datasets X and Y, in our case between RS-fMRI connectivity measures and clinical symptoms.

#### 2.5.1 Permutation test

Traditionally, the significance of canonical correlations is established using a Wilk’s lambda statistic and this was also used in the Drysdale study (Conor Liston, personal communication). This statistic has an approximately *chi-square* null distribution with *pq* degrees of freedom, *p* and *q* being the number of variables in X and Y. This significance test, however, does not take into account the feature pre-selection step that selected RS-fMRI connectivity features most correlated with clinical symptoms. As this pre-selection step was done in the same dataset as the CCA was performed on and tested, this likely results in too optimistic p-values. To avoid this issue, we performed a permutation test of the whole procedure, including feature selection followed by CCA. The whole feature selection and CCA cycle was repeated for each permutation with the rows of clinical symptoms shuffled so that they no longer corresponded to rows of RS-fMRI connectivity features. We performed 499 permutations, which created a null distribution of canonical correlations and estimated the Wilk’s lambda statistic. The interpretation of Wilk’s lambda statistic is also important, because it does not describe the significance of a single component in isolation. Instead, the first canonical correlation is defined as det(E)/det(E +H), where E is the error sum of squares and cross products matrix and H is model sum of squares and cross products matrix. Its significance should be interpreted as the significance of the whole decomposition, not the first component. The significance of the Wilk’s lambda statistic for the second canonical correlation is interpreted as the significance of the whole decomposition after removing the variance accounted for by the first canonical correlation and so forth. In addition, if the canonical correlation from a given model order (e.g. first canonical correlation) is not significant, all correlations of a lower order (e.g. second onwards) should not be taken to be significant either, even if one or more of the derived p-values show nominal significance [17].

#### 2.5.2 Cross-validation

CCA is prone to overfitting and although canonical correlations may seem high and even be statistically significant, they are often much lower in an independent dataset (see e.g. [8]). This might give an impression that the found association between modalities (RS-fMRI connectivity measures and clinical symptoms) is much stronger than it would be in an independent hold-out dataset. In the original study, the canonical correlation in the independent data set was not evaluated directly in an independent dataset, but rather the authors relied on the derived biotypes to have similar symptom profiles in the independent evaluation dataset.

Here we chose to estimate the magnitude of the canonical correlation in independent datasets directly, using a stratified 10-fold cross-validation. The dataset was divided into ten subsets with an approximately constant number of subjects from each scan location across all subsets. Nine subsets were used as a training set and the remaining subset as a test set. A feature selection procedure, as described above, was performed using subjects from the training set only. In the test set, canonical variates and their respective canonical correlations were created using coefficients from the CCA performed in the training set.

#### 2.5.3 Stability of canonical loadings

Since CCA frequently yields unstable solutions, we also examined the stability of canonical loadings (i.e. structure coefficients, a univariate correlation between a variable and canonical variate) [17] under resampling of the data. We repeated the whole feature selection and CCA procedure multiple times always with leaving one subject out of the analysis. This produces a distribution of canonical loadings and thus allows us to estimate their stability, and therefore uncertainty, under small perturbations of the data (here by exchanging one subject) taking into an account both the feature selection step and the CCA step.

### 2.6 Clustering analysis

In the original study, the first two canonical variates of the RS-fMRI connectivity features were used as input for the clustering analysis. The underlying idea was to constrain the clustering analysis to a low dimensional representation of brain connectivity features that are clinically relevant. To make our analysis comparable to the original study, we decided to perform a clustering analysis on two different sets of canonical variates. First, as in the original study, we performed clustering analysis on the first two RS-fMRI connectivity canonical variates, which were the two RS-fMRI components with the highest canonical correlations. Second, because the first two components in our analysis may not correspond to the first two components identified in the original study, we visually selected two canonical variates that showed the most similar clinical profiles to those identified in the original study.

Then, we performed the same hierarchical clustering procedure as in the Drysdale et al. study, i.e., using the Euclidean distance measure and Ward’s D linkage method, which minimize the total within-cluster variance. As a measure of quality of our clustering solution, we computed the CH index as in the original study (variance ratio criterion or Calinski-Harabasz index), which is the ratio of between-cluster variance and within-cluster variance. As an additional metric, we also compared model order using the silhouette index, which compares average within cluster distances to average distances between points from different clusters.

In the original study, the decision to identify four clusters as the best clustering solution was made partly because the CH index was maximized for the four cluster solution. This, by itself, is not a statistical test or evidence of the existence of 4 clusters. Specifically, we don’t know if the derived CH index was significantly higher than what would have been expected under the null hypothesis of data with no underlying clusters. Here we devise a procedure, similar to the one proposed in [18], to test the statistical significance of the observed CH index. In this procedure, the null hypothesis is that the data came from a single 2-dimensional Gaussian distribution (i.e. distribution with no underlying clusters). Specifically, first, we estimated a covariance matrix between the two canonical variates used for the clustering analysis. Second, we repeatedly took random samples of the size of our dataset (187) from a bivariate Gaussian distribution defined by this covariance matrix. Third, we ran the same hierarchical clustering procedure as we performed on the real data on each random sample and calculated the best obtained CH and silhouette index, thus obtaining an empirical null distribution of these indices. The p-value was then defined as a proportion of the calculated indices in the null distribution smaller than what we observed in the real data.

#### 2.6.1 Stability of clustering

To reliably interpret the derived clusters, it is important to evaluate if the clustering assignment is stable under small perturbations to the data. In other words, if the same procedure was repeated using a similar dataset, would we identify similar clusters and would we assign the same subjects to the same clusters? This is a different question than if the clusters are statistically significant, because cluster assignment might be stable even if there are no real clusters in the data. On the other hand, it is possible to obtain clearly distinct clusters that are unreliable and cannot be reproduced in a different dataset. For this, we employed the same leave-one-out procedure as for the estimation of the stability of the canonical loadings. The whole feature selection, CCA, and hierarchical clustering procedure were repeated for each subject, always with one subject left out of the training process. This allowed us to estimate the stability of the canonical variates under slight perturbation of the data and subsequently the stability of the whole clustering procedure that is based on these canonical variates.

### 2.7 Code availability

The code used to perform data analysis can be found in supplementary materials.

## 3. RESULTS

### 3.1 Sample characteristics

Sample characteristics are provided in Table 2.

**Table 2:** Sample characteristics for the NESDA and MOTAR samples

| Study | N | Age | Age SD | N Female | % Female | MDD | Anxiety | Comorbid |
| --- | --- | --- | --- | --- | --- | --- | --- | --- |
| **NESDA** | 151 | 36.42 | 10.92 | 101 | 67 | 53 | 45 | 53 |
| **MOTAR** | 36 | 36.69 | 12.37 | 23 | 66 | 10 | 2 | 24 |
| **Total** | 187 | 36.48 | 10.93 | 124 | 66 | 63 | 47 | 77 |

### 3.2 CCA significance

Canonical correlations were 0.99 and 0.98 for the first two pairs of canonical variates. As can be seen in the null distribution provided in Figure 2, canonical correlations this high are not unusual even if there is no actual correspondence between X and Y (as determined by a permutation test). Indeed, the respective p-values of the permutation tests were not significant (p=0.06 and p=0.90), neither were they significant according to the Wilk’s lambda statistics (p=0.6, and p=0.8), which measured the significance of the whole decomposition. Our permutation testing procedure took into account that connectivity features had been selected based on their correlation with clinical features. Because the Drysdale et al. study did not test the significance of their CCA solution in the same way, it remains to be confirmed whether the canonical variates identified in their original study were significant, although authors did provide indirect evidence of this using an independent validation sample.

**Figure 2:**
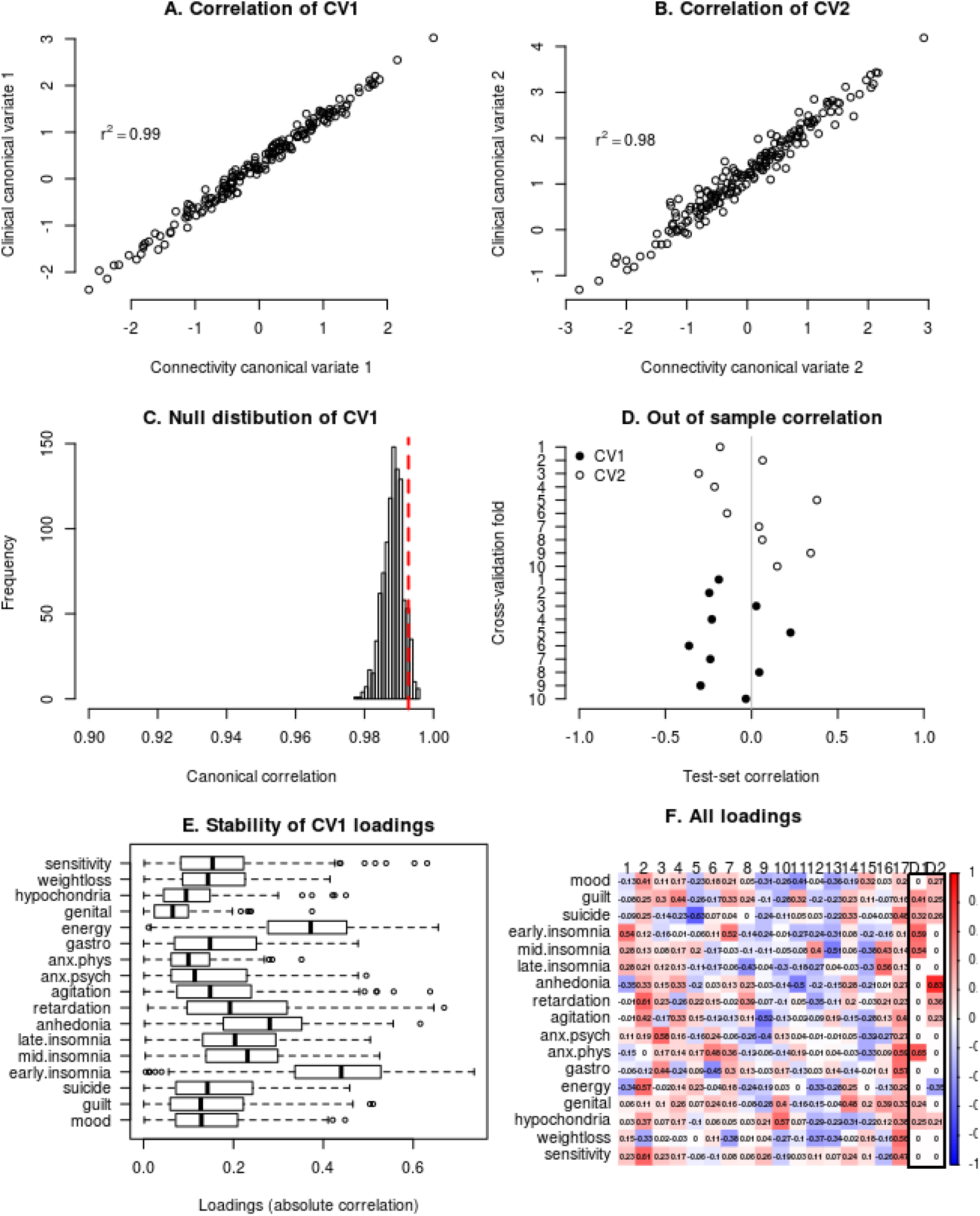
**A, B)** CCA finds a linear combination (canonical variate) of brain connectivity features that maximizes correlation with a linear combination of clinical symptoms. Canonical correlations are high and comparable to the original study (0.95 and 0.91). **C)** The null distribution of the first canonical correlation obtained using permutation test. Although canonical correlations in A and B are seemingly high, they are also high under the null hypothesis and thus not statistically significant. **D)** Out of sample canonical correlation for first two canonical pairs estimated by 10 fold cross-validation. Each point represents out of sample canonical correlation for each cross-validation fold. Although the canonical correlation was high in the training set as showed in A and B, id dropped to a chance level correlation in the test sets. **E)** Canonical loadings for the first canonical variate and their stability under resampling of the data using leave-one-out (jack-knife) procedure. **F)** Clinical canonical loadings for all canonical variates (1-17) and first two reported in the original study (D1-D2)

#### 3.2.1 Out of sample canonical correlation

Using 10-fold cross-validation, the average out of sample canonical correlation was −0.03 and −0.3 for the first and second canonical pair respectively. This shows that even a high within-sample canonical correlation might not necessarily be reproducible in an independent test set. The original study used an independent evaluation set of 92 subjects. However, the authors did not perform a CCA analysis in the independent sample directly and thus did not provide canonical correlations for the independent validation dataset. Instead, they demonstrated that subjects assigned to clusters in the validation set according to their RS-fMRI connectivity features showed similar clinical profiles as the clusters identified in the training set.

#### 3.2.2 CCA similarity of loadings

A side-by-side comparison of canonical loadings (univariate correlation between each variable and the canonical variate) of all our resulting canonical variates and the first two canonical variates reported in Drysdale et al. are provided in Figure 2f. We also conducted an analysis of stability of these loadings under small resampling of the data by repeating the feature selection and CCA procedure 187 times, each time without one subject left out of the analysis. The results for this analysis can be seen in Figure 2e. It can be seen that even by changing one subject in the pipeline, individual loadings changed dramatically. Because of this instability and the fact that our canonical variates were not statistically significant, it is not meaningful to compare our loadings with loadings found in the original study.

### 3.3 Clustering significance

In our dataset, a 4-cluster solution showed the highest CH (110) and 2 cluster solution the highest silhouette index (0.33). However, using a simulation approach described in the methods section, these indices were not statistically significant (p=0.36 and p=0.71 for CH and silhouette index, respectively). That means that it is not unusual to observe such high CH and silhouette indices, even when the hierarchical clustering is performed on a normally distributed data set (data with no clusters). Formally, this means that we cannot reject the null hypothesis of the data coming from a single Gaussian distribution (Figure 3). In the original study, this was not tested. Therefore, we cannot say if the data in the original study really formed clusters, instead of just random fluctuation of the data.

**Figure 3:**
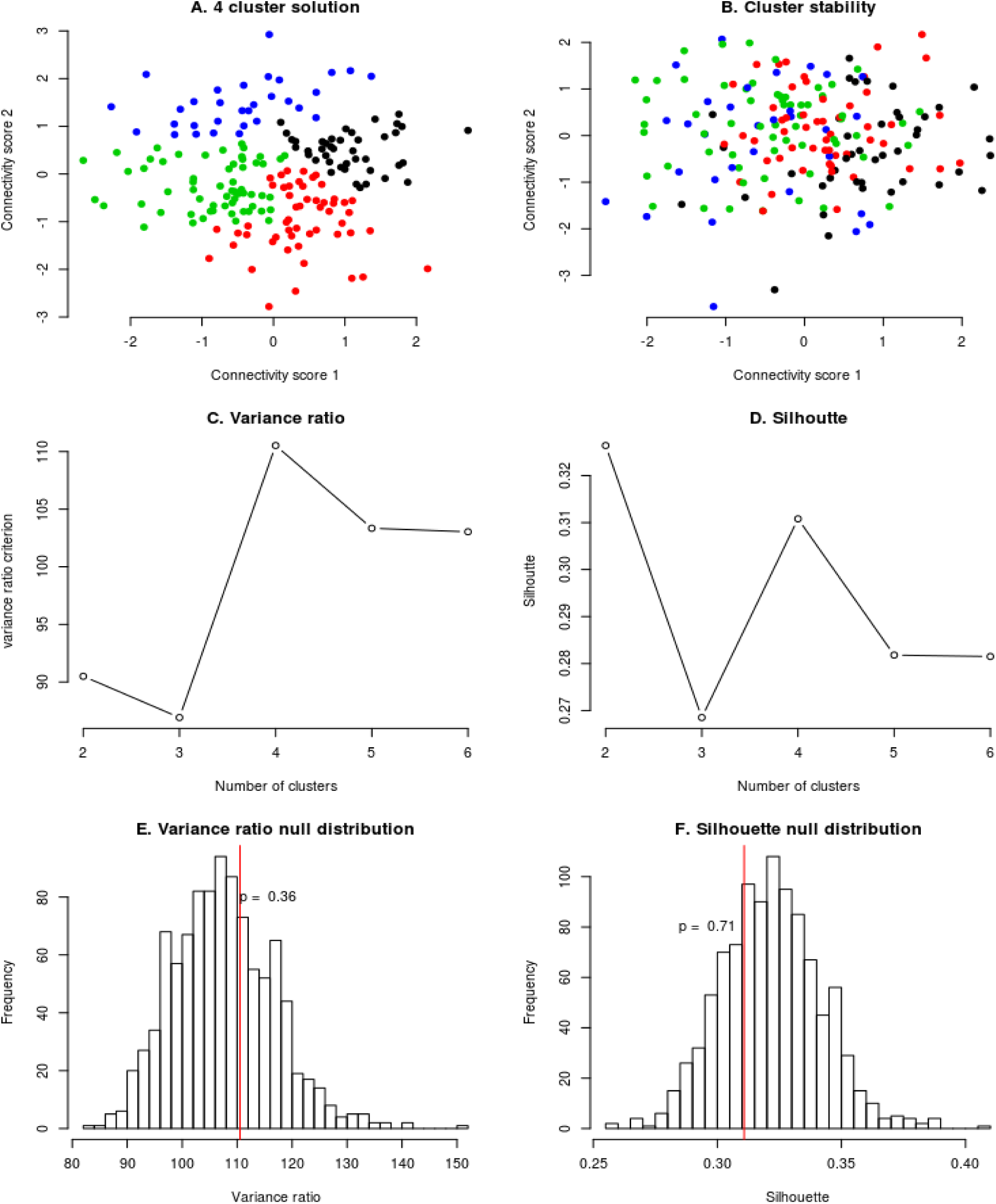
**A)** obtained 4-cluster solution using hierarchical clustering. **B)** Stability of a cluster assignment. Each subject is shown with the same color as it had in A, but the connectivity scores are recomputed under a small perturbation of the data - leaving one subject out of the feature selection and CCA procedure. **C)** Variance ratio criterion is maximized at 4 clusters as in the original study. **D)** Silhouette index is maximized at 2 clusters. **E, F)** Null distribution of Variance ratio and silhouette indices. Showing that although these indices are maximized at 4 and 2 clusters respectively, these results are not unusual even for the data simulated from a distribution with no clusters. Therefore these criteria do not imply evidence for the existence of clusters in our data or in the data presented in the original study.

#### 3.3.1 Cluster stability

We evaluated cluster assignment stability under slight resampling of the data by performing the feature selection and CCA procedure again, with one subject left out of the pipeline. The results are shown in the Figure 3a and 3b. The figures show the position of each subject with respect to the first two canonical variates and therefore how clusters changed just by changing one subject in the pipeline. Stability of cluster assignment was performed in the original study by bootstrap procedure after feature selection and CCA, thus not taking instability of these two steps into account.

#### 3.3.2 Clustering similarity

Since we did not find evidence for clusters in our data, and the cluster assignment was not stable, we did not consider it meaningful to describe clusters in terms of their symptom profiles or compare the clusters in our study to clusters identified in the original study.

## 4. DISCUSSION

Here, we performed a partial replication of the study of the study by Drysdale and colleagues [1] using a completely independent dataset approximately matching the original study in terms of sample size and derived measures. We followed the analysis steps of the original study as closely as possible and found similar results in terms of the magnitude of canonical correlations and number of clusters. However, after performing additional tests that were not performed in the original study, which took into account that the RS-fMRI connectivity features were selected based on their correlation with clinical features before performing CCA on these connectivity and clinical features, even the high canonical correlations that we observed were not statistically significant and they did not replicate outside of the training set. By using the same criteria for selecting the number of clusters as in the original study, we found an optimal four cluster solution. However, we showed that this cluster solution would happen even if the data came only from a single Gaussian distribution with no underlying clusters.

### 4.1 Statistical significance of canonical correlations

The first two canonical correlations between brain connectivity and clinical symptom measures that we observed in our data were high (0.99 and 0.98). However, they were not statistically significant as determined by permutation testing. In addition, using cross-validation, the canonical correlations dropped to approximately 0 in the test set. This is not unexpected because of the high number of variables included compared to the number of subjects in this sample, which leads to severe overfitting. CCA is known to be unstable; for example, introductory texts recommend between 10 to 42 subjects per variable in order to obtain a reliable CCA model [19,20], but the pipeline used in the original study that we followed had around 1.3 subjects per variable. Another important contribution for the overly optimistic canonical correlations is the initial feature selection step that selected 150 connectivity features in our study (178 in the original study) out of ∼30,000 brain connectivity measures that were most correlated with the clinical symptoms in the same dataset in which the CCA was performed.

In the original study by Drysdale and colleagues, this feature selection step was not taken into account when estimating the statistical significance of the canonical correlations, thus the reported p-values were likely inflated. Moreover, the replication of canonical correlations out of sample was not shown directly in the study by Drysdale and colleagues. Despite this, the authors did provide indirect evidence for a reliable relationship between brain connectivity measures and clinical symptoms in a subset of subjects left out completely from the primary analysis (training set). These subjects were assigned to clusters based on a support vector machine classifier using only their connectivity features as predictors, and these clusters had similar clinical symptom profiles in their cluster discovery and replication sets, which would not be possible if the canonical correlations were spurious.

### 4.2 Similarity of canonical loadings

Due to the overfitting of CCA discussed above, the canonical loadings we obtained were unstable, which makes their comparison with loadings reported in the original study difficult and unreliable. Despite that, loadings of our fourth canonical variate were most similar to the loadings of the second canonical variate reported in the original study by Drysdale and colleagues. However, our canonical variates were not statistically significant, therefore this similarity cannot be interpreted as a replication of the same biological to clinical connection as found in the original study.

### 4.3 Clustering analysis

A problem with many clustering algorithms that is not commonly recognized is that they always yield clusters, regardless of the structure of the data, even if there are no clusters at all [18]. Many procedures employed to determine the optimal number of clusters, including the one used in the original study, are therefore more heuristic and do not provide a statistical test of the underlying structure of the data. In the original study, a four-cluster solution was decided to be optimal mainly because the CH criterion, a specific numerical value describing how well the data form clusters, was maximized by four clusters. According to this criterion, we would have chosen an optimal number of clusters to be four (or two according to the silhouette criterion) in our data. However, after a closer examination, we observed that a CH index as high or higher as we observed, can be easily obtained just by running the same hierarchical clustering procedure on data randomly sampled from a distribution that does not contain any clusters (in this case Gaussian distribution). Or in other words, according to the CH index, we could not reject the null hypothesis that our data came from a single Gaussian distribution. No test for the existence of clusters was performed in the original study and the presented data in Figure 1f of the original study looked more like a continuous distribution instead of 4 clusters. Therefore, we think that the original study provided insufficient evidence to conclude the existence of any number of distinct “biotypes” of depression.

Although the absence of clusters would change the conclusion of the original study, it is not necessarily detrimental to the significance of the results. The found biological axes related to different depressive symptoms are important in their own merit, without subsequent arbitrary dichotomization into four biotypes. Two canonical variates already provide a parsimonious representation of the data and dividing them further into four subtypes would not provide any more insights into mechanisms of depression (especially if these subtypes are spurious).

Assuming biotypes is also detrimental for the sake of clinical utility, such as predicting the probability of a TMS treatment response. Since the subtypes were predictive of TMS treatment response and were based on the underlying canonical variates, it is reasonable to assume that the probability of response varies smoothly with respect to the canonical variates. Using only discrete subtypes for prediction assumes that all the subjects in one “biotype” have the same chance for response. Also, very similar subjects might get significantly different predictions. If a subject would move slightly from biotype 1 to bordering biotype 2, his predicted TMS response chance would jump from 80% to 20%.

On the other hand, using subject-specific connectivity scores alone, without additional arbitrary dichotomization into “biotypes,” would allow making an individualized prediction for each individual, in line with goals of personalized precision medicine. A clinical decision can then be made for each patient individually according to their treatment response probability instead of the average treatment response probability of the whole group (e.g. biotype). Critically, the availability of quantitative measures means that cut-off points for various levels of severity can be changed and fine-tuned as more data from future studies become available — as has been done for diseases such as hypertension. Severity cut-points explicitly acknowledge dimensions and move away from traditional single disorder models. Such a dimensional approach, which captures the full spectrum of brain connectivity alterations, provides an empirical and coherent framework to accommodate comorbidity and sub-threshold symptoms.

Due to the clinical heterogeneity of many psychiatric disorders and the quest for personalized medicine, there is a tendency towards subtyping and expanding psychiatric nosology. However, the presumption of distinct and homogeneous subtypes might not be clinically useful and might not represent the underlying biology. Many clustering approaches will always produce some clusters and would so even for uniformly distributed data. It is, therefore, crucial to distinguish real biologically or clinically meaningful subtypes from random fluctuation of the data. This is not an easy task, however, several methods exist. One possibility is to use simulations to create an empirical null distribution of clustering statistics, similar to what we used here as proposed by [18] or use model-based approaches, such as latent class analysis or Gaussian mixture models, where the model fit can be tested directly.

### 4.4 Recommendations for future studies

We have several recommendations for future studies. First, to avoid overfitting and unstable results of CCA, we advise to either reduce the number of features by using a dimensionality reduction method, such as PCA, ICA or factor analysis as used in [7], or to use a regularized version of CCA, or both as used in [8]. Second, if a feature selection step is involved, it is necessary to take this into account in the statistical testing procedure, either by doing a statistical test in an independent test set or by incorporating this selection step into a permutation testing procedure [21]. For clustering analysis, it is necessary first to answer the question if there are actually real clusters in the data or just random fluctuations. Clustering coefficients and cluster assignment stability evaluation do not test for this. To estimate cluster stability assignment it is important to take the whole procedure into account, including feature selection and CCA, which might show that even seemingly stable clusters are unstable. Finally, if the goal is a clinically useful prediction and high accuracy, continuous variables should be preferred before dichotomizing data into clusters because they contain more information and thus lead to better prediction.

### 4.5 Limitations

A limitation of this replication attempt is that our sample characteristics were different from the original study, which included only subjects with a currently active episode of a treatment-resistant major depressive disorder. In contrast, we included a wider range of subjects with mild to severe symptoms, recruited from the general populations, primary and secondary care, as well as a wider range of diagnoses including MDD, anxiety and comorbid MDD with anxiety. However, this could also be considered a strength, because a wider range of symptoms should also translate to more diverse biology and thus make the connection between clinical symptoms to biology more apparent. Another limitation is that we used a different clinical measurement tool (IDS instead of HAM-D), however, we matched the items as closely as possible. Lastly, we did not replicate all parts of the original study, such as prediction of treatment response or classification of depressed and healthy subjects.

### 4.6 Conclusion

To the best of our knowledge, this is the first attempt to replicate the important findings relating clinical symptoms and biological subtypes reported in the Drysdale et al. study. Using additional statistical procedures, the method used in the original study did not provide stable or statistically significant biotypes of depression and anxiety in our independent dataset. Furthermore, we have argued that the evidence for the existence of 4 distinct biotypes presented in the original study is not convincingly demonstrated and the results should be interpreted with care. However, even without partitioning patients into biotypes, the existence of continuous biological axes related to symptoms of depression might be even more useful for our understanding of biological mechanisms of depression and clinical practice. However, unfortunately, we were not able to replicate such dimensional axes in our data.

## Acknowledgements

This work was supported by the Geestkracht program of the Netherlands Organization for Health Research and Development (Zon-Mw, grant number 10-000-1002) and is also supported by participating universities and mental health care organizations (VU University Medical Center, GGZ inGeest, Arkin, Leiden University Medical Center, GGZ Rivierduinen, University Medical Center Groningen, Lentis, GGZ Friesland, GGZ Drenthe, Institute for Quality of Health Care (IQ Healthcare), Netherlands Institute for Health Services Research (NIVEL) and Netherlands Institute of Mental Health and Addiction (Trimbos). This work was also supported by Neuroscience Amsterdam (PoC-2014-NMH-02). LS and RD are supported by The Netherlands Brain Foundation Grant number (F2014 (1)-24). AM gratefully acknowledges support from the NWO under a VIDI fellowship (grant number 016.156.415). MJvT was supported by a VENI grant (NWO grant number 016.156.077). We also gratefully acknowledge the assistance from Dr. Conor Liston for providing us with additional details necessary to replicate the analysis pipeline for Drysdale et al 2017.

